# Integrating Hi-C links with assembly graphs for chromosome-scale assembly

**DOI:** 10.1101/261149

**Authors:** Jay Ghurye, Arang Rhie, Brian P. Walenz, Anthony Schmitt, Siddarth Selvaraj, Mihai Pop, Adam M. Phillippy, Sergey Koren

## Abstract

Long-read sequencing and novel long-range assays have revolutionized *de novo* genome assembly by automating the reconstruction of reference-quality genomes. In particular, Hi-C sequencing is becoming an economical method for generating chromosome-scale scaffolds. Despite its increasing popularity, there are limited open-source tools available. Errors, particularly inversions and fusions across chromosomes, remain higher than alternate scaffolding technologies. We present a novel open-source Hi-C scaffolder that does not require an *a priori* estimate of chromosome number and minimizes errors by scaffolding with the assistance of an assembly graph. We demonstrate higher accuracy than the state-of-the-art methods across a variety of Hi-C library preparations and input assembly sizes. The Python and C++ code for our method is openly available at https://github.com/machinegun/SALSA

**Author summary:** Hi-C technology was originally proposed to study the 3D organization of a genome. Recently, it has also been applied to assemble large eukaryotic genomes into chromosome-scale scaffolds. Despite this, there are few open source methods to generate these assemblies. Existing methods are also prone to small inversion errors due to noise in the Hi-C data. In this work, we address these challenges and develop a method, named SALSA2. SALSA2 uses sequence overlap information from an assembly graph to correct inversion errors and provide accurate chromosome-scale assemblies.

## Introduction

Genome assembly is the process of reconstructing a complete genome sequence from significantly shorter sequencing reads. Most genome projects rely on whole genome shotgun sequencing which yields an oversampling of each genomic locus. Reads originating from the same locus are identified using assembly software, which can use these overlaps to reconstruct the genome sequence [1, 2]. Most approaches are based on either a de Bruijn [3] or a string graph [4] formulation. Repetitive sequences exceeding the sequencing read length [5] introduce ambiguity and prevent complete reconstruction. Unambiguous reconstructions of the sequence are output as “unitigs” (or often “contigs”). Ambiguous reconstructions are output as edges linking unitigs. Scaffolding utilizes long-range linking information such as BAC or fosmid clones [6, 7], optical maps [8–10], linked reads [11–13], or chromosomal conformation capture [14] to order and orient unitigs. If the linking information spans large distances on the chromosome, the resulting scaffolds can span entire chromosomes or chromosome arms.

Hi-C is a sequencing-based assay originally designed to interrogate the 3D structure of the genome inside a cell nucleus by measuring the contact frequency between all pairs of loci in the genome [15]. The contact frequency between a pair of loci strongly correlates with the one-dimensional distance between them. Hi-C data can provide linkage information across a variety of length scales, spanning tens of megabases. As a result, Hi-C data can be used for genome scaffolding. Shortly after its introduction, Hi-C was used to generate chromosome-scale scaffolds [16–20].

LACHESIS [16] is an early method for Hi-C scaffolding which first clusters unitigs into a user-specified number of chromosome groups and then orients and orders the unitigs in each group independently to generate scaffolds. Thus, the scaffolds inherit any assembly errors present in the unitigs. The original SALSA1 [21] method first corrects the input assembly, using a lack of Hi-C coverage as evidence of error. It then orients and orders the corrected unitigs to generate scaffolds. Recently, the 3D-DNA [20] method was introduced and demonstrated on a draft assembly of the *Aedes aegypti* genome. 3D-DNA also corrects the errors in the input assembly and then iteratively orients and orders unitigs into a single megascaffold. This megascaffold is then broken, identifying chromosomal ends based the on Hi-C contact map.

There are several shortcomings common across currently available tools. They are sensitive to input assembly contiguity and Hi-C library variations and require tuning of parameters for each dataset. Inversions are common when the input unitigs are short, as orientation is determined by maximizing the interaction frequency between unitig ends across all possible orientations [16]. When unitigs are long, there are few interactions spanning the full length of the unitig, making the true orientation apparent from the higher weight of links. However, in the case of short unitigs, there are interactions spanning the full length of the unitig, making the true orientation have a similar weight to incorrect orientations. Biological factors, such as topologically associated domains (TADs) also confound this analysis [22].

SALSA1 [21], addressed some of these challenges, such as not requiring the expected number of chromosomes beforehand and correcting assemblies before scaffolding them with Hi-C data. We showed that SALSA1 worked better than the most widely used method, LACHESIS [16]. However, SALSA1 often did not generate chromosome-sized scaffolds. The contiguity and correctness of the scaffolds depended on the coverage of Hi-C data and required manual data-dependent parameter tuning. Building on this work, SALSA2 does not require manual parameter tuning and is able to utilize all the contact information from the Hi-C data to generate near optimal sized scaffolds permitted by the data using a novel iterative scaffolding method. In addition to this, SALSA2 enables the use of an assembly graph to guide scaffolding, thereby minimizing errors, particularly orientation errors.

In this work, we introduce SALSA2 - an open source software that combines Hi-C linkage information with the ambiguous-edge information from a genome assembly graph to better resolve unitig orientations. We also propose a novel stopping condition, which does not require an *a priori* estimate of chromosome count, as it naturally stops when the Hi-C information is exhausted. We show that SALSA2 has fewer orientation, ordering, and chimeric errors across a wide range of assembly contiguities. We also demonstrate robustness to different Hi-C libraries with varying intra-chromosomal contact frequencies. When compared to 3D-DNA, SALSA2 generates more accurate scaffolds across most conditions. To our knowledge, this is the first method to leverage assembly graph information for scaffolding Hi-C data.

## 1 Methods

Figure 1(A) shows the overview of the SALSA2 pipeline. SALSA2 begins with a draft assembly generated from long reads such as Pacific Biosciences [23] or Oxford Nanopore [24]. SALSA2 requires the unitig sequences and, optionally, a GFA-formatted assembly graph [25] representing the ambiguous reconstructions. Hi-C reads are aligned to the unitig sequences, and unitigs are optionally split in regions lacking Hi-C coverage. A hybrid scaffold graph is constructed using both ambiguous edges from the GFA and edges from the Hi-C reads, scoring edges according to a “best buddy” scheme. Scaffolds are iteratively constructed from this graph using a greedy weighted maximum matching. A mis-join detection step is performed after each iteration to check if any of the joins made during this round are incorrect. Incorrect joins are broken and the edges blacklisted during subsequent iterations. This process continues until the majority of joins made in the prior iteration are incorrect. This provides a natural stopping condition, when accurate Hi-C links have been exhausted. Below, we describe each of the steps in detail.

**Fig 1.**
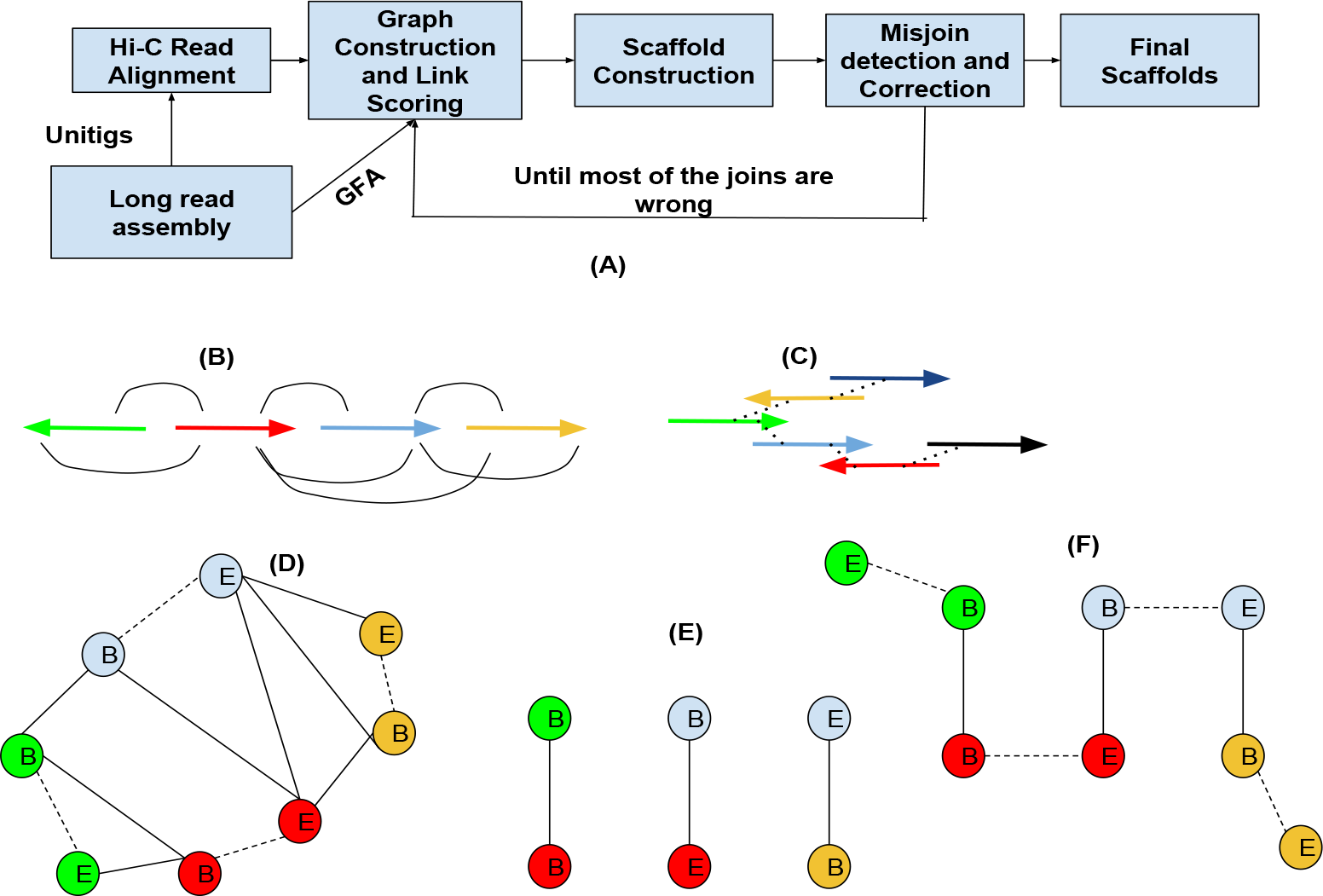
(A) Overview of the SALSA2 scaffolding algorithm. (B) Linkage information obtained from the alignment of Hi-C reads to the assembly. (C) The assembly graph obtained from the assembler. (D) A hybrid scaffold graph constructed from the links obtained from the Hi-C read alignments and the overlap graph. Solid edges indicate the linkages between different unitigs and dotted edges indicate the links between the ends of the same unitig. (E) Maximal matching obtained from the graph using a greedy weighted maximum matching algorithm. (F) Edges between the ends of same unitigs are added back to the matching.

### 1.1 Hi-C library preparation

Hi-C methods first crosslink a sample (cells or tissues) to preserve the genome conformation. The crosslinked DNA is then digested using restriction enzymes (in this case GATC and GANTC). The single-stranded 5’-overhangs are then filled in causing digested ends to be labeled with a biotinylated nucleotide. Next, spatially proximal digested ends of DNA are ligated, preserving both short- and long-range DNA contiguity. The DNA is then purified and sheared to a size appropriate for Illumina short-read sequencing. After shearing, the biotinylated fragments are enriched to assure that only fragments originating from ligation events are sequenced in paired-end mode via Illumina sequencers to inform DNA contiguity.

### 1.2 Read alignment

Hi-C paired end reads are aligned to unitigs using the BWA aligner [26](parameters: -t 12 -B 8) as single end reads. First, the reads mapping at multiple locations are ignored as they can cause ambiguities while scaffolding. Reads which align across ligation junctions are chimeric and are trimmed to retain only the start of the read which aligns prior to the ligation junction. After filtering the chimeric reads, the pairing information is restored. Any PCR duplicates in the paired-end alignments are removed using Picard tools [27]. Read pairs aligned to different unitigs are used to construct the initial scaffold graph. The suggested mapping pipeline is available at http://github.com/ArimaGenomics/mapping_pipeline.

### 1.3 Unitig correction

As any assembly is likely to contain mis-assembled sequences, SALSA2 uses the physical coverage of Hi-C pairs to identify suspicious regions and break the sequence at the likely point of mis-assembly. We define the physical coverage of a Hi-C read pair as the region on the unitig spanned by the start of the leftmost fragment and the end of the rightmost fragment. A drop in physical coverage indicates a likely assembly error. In SALSA1, unitigs are split when a fixed minimum coverage threshold is not met. A drawback of this approach is that coverage can vary, both due to sequencing depth and variation in Hi-C link density.

Figure 2 sketches the new unitig correction algorithm implemented in SALSA2. Instead of the single coverage threshold used in SALSA1, a set of suspicious intervals is found with a sweep of thresholds. For different thresholds for low coverage cutoffs, we find the continuous stretches of regions which have lower physical coverage than the cutoff under consideration. We find such intervals of continuous low coverage for different cutoffs. Using the collection of intervals as an interval graph, we find the maximal clique. This maximal clique represents the region of the unitig which had low coverage for all the cutoffs. This can be done in *O*(*NlogN*) time, where *N* is the number of intervals. For a maximal clique, the region between the start and end of the smallest interval in the clique is flagged as a mis-assembly and the unitig is split into three pieces — the sequence to the left of the region, the junction region itself, and the sequence to the right of the region. The intuition behind choosing the smallest interval is to accurately pinpoint the location of assembly error.

**Fig 2.**
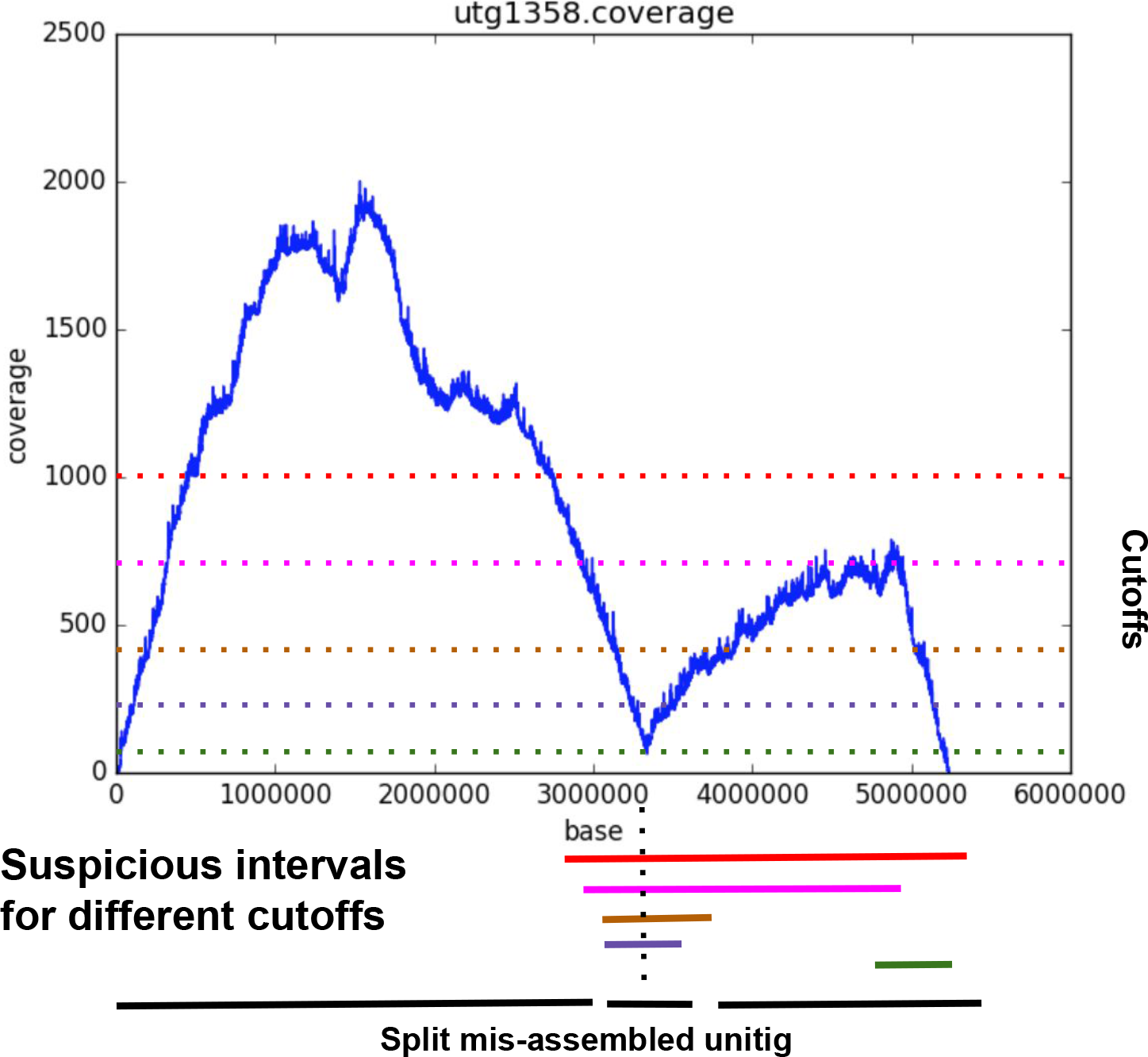
Example of the mis-assembly detection algorithm in SALSA2. The plot shows the position on x-axis and the physical coverage on the y-axis. The dotted horizontal lines show the different thresholds tested to find low physical coverage intervals. The lines at the bottom show the suspicious intervals identified by the algorithm. The dotted line through the intervals shows the maximal clique. The smallest interval (purple) in the clique is identified as mis-assembly and the unitig is broken in three parts at its boundaries.

### 1.4 Assembly graph construction

For our experiments, we use the unitig assembly graph produced by Canu [28] (Figure 1(C)), as this is the more conservative graph output. SALSA2 requires only a GFA format [25] representation of the assembly. Since most long read genome assemblers such as FALCON [29], miniasm [25], Canu [28], and Flye [30] provide assembly graphs in GFA format, their output is compatible with SALSA2 for scaffolding.

### 1.5 Scaffold graph construction

The scaffold graph is defined as *G*(*V*, *E*), where nodes *V* are the ends of unitigs and edges *E* are derived from the Hi-C read mapping (Figure 1B). The idea of using unitig ends as nodes is similar to that used by the string graph formulation [4].

Modeling each unitig as two nodes allows a pair of unitigs to have multiple edges in any of the four possible orientations (forward-forward, forward-reverse, reverse-forward, and reverse-reverse). The graph then contains two edge types — one explicitly connects two different unitigs based on Hi-C data, while the other implicitly connects the two ends of the same unitig.

As in SALSA1, we normalize the Hi-C read counts by the frequency of restriction enzyme cut sites in each unitig. This normalization reduces the bias in the number of shared read pairs due to the unitig length as the number of Hi-C reads sequenced from a particular region are proportional to the number of restriction enzyme cut sites in that region. For each unitig, we denote the number of times a cut site appears as *C*(*V*). We define edges weights of *G* as:

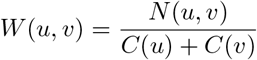

where *N* (*u*, *v*) is the number of Hi-C read pairs mapped to the ends of the unitigs *u* and *v*.

We observed that the globally highest edge weight does not always capture the correct orientation and ordering information due to variations in Hi-C interaction frequencies within a genome. To address this, we defined a modified edge ratio, similar to the one described in [20], which captures the relative weights of all the neighboring edges for a particular node.

The best buddy weight *BB*(*u*, *v*) is the weight *W* (*u*, *v*) divided by the maximal weight of any edge incident upon nodes *u* or *v*, excluding the (*u*, *v*) edge itself. Computing best buddy weight naively would take *O*(|*E*|^2^) time. This is computationally prohibitive since the graph, *G*, is usually dense. If the maximum weighted edge incident on each node is stored with the node, the running time for the computation becomes *O*(|*E*|). We retain only edges where *BB*(*u*, *v*) > 1. This keeps only the edges which are the best incident edge on both *u* and *v*. Once used, the edges are removed from subsequent iterations. Thus, the most confident edges are used first but initially low scoring edges can become best in subsequent iterations.

For the assembly graph, we define a similar ratio. Since the edge weights are optional in the GFA specification and do not directly relate to the proximity of two unitigs on the chromosome, we use the graph topology to establish this relationship. Let 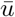 denote the reverse complement of the unitig *u*. Let *σ*(*u*, *v*) denote the length of shortest path between *u* and *v*. For each edge (*u*, *v*) in the scaffold graph, we find the shortest path between unitigs *u* and *v* in every possible orientation, that is, *σ*(*u*, *v*), 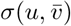, 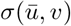 and 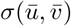. With this, the score for a pair of unitigs is defined as follows:

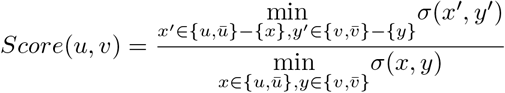

where *x* and *y* are the orientations in which *u* and *v* are connected by a shortest path in the assembly graph. Essentially, *Score*(*u*, *v*) is the ratio of the length of the second shortest path to the length of the shortest path in all possible orientations. Once again, we retain edges where *Score*(*u*, *v*) > 1. If the orientation implied by the assembly graph differs from the orientation implied by the Hi-C data, we remove the Hi-C edge and retain the assembly graph edge (Figure 1D). Computing the score graph requires |*E*| shortest path queries, yielding total runtime of *O*(|*E*| (|*V*| + |*E*|)) since we do not use the edge weights.

### 1.6 Unitig layout

Once we have the hybrid graph, we lay out the unitigs to generate scaffolds. Since there are implicit edges in the graph *G* between the beginning and end of each unitig, the problem of computing a scaffold layout can be modeled as finding a weighted maximum matching in a general graph, with edge weights being our ratio weights. If we find the weighted maximum matching of the non-implicit edges (that is, edges between different unitigs) in the graph, adding the implicit edges to this matching would yield a complete traversal. However, adding implicit edges to the matching can introduce a cycle. Such cycles are prevented by removing the lowest weight non-implicit edge. Computing a maximal matching takes *O*(|*E*| |*V*|^2^) time [31]. We iteratively find a maximum matching in the graph by removing nodes found in the previous iteration. Using the optimal maximum matching algorithm this would take *O*(|*E*‖*V*|^3^) time, which would be extremely slow for large graphs. Instead, we use a greedy maximal matching algorithm which is guaranteed to find a matching within 1/2-approximation of the optimum [32]. The greedy matching algorithm takes *O*(|*E*|) time, thereby making the total runtime *O*(|*V*‖*E*|). The algorithm for unitig layout is sketched in Algorithm 1. Figure 1(D - F) show the layout on an example graph. Contigs which were not scaffolded are inserted in the large scaffolds with the method used in SALSA1.

#### Algorithm 1 Unitig Layout Algorithm

~~~
  *E* : Edges sorted by the best buddy weight
  *M* : Set to store maximal matchings
  *G* : The scaffold graph
  **while** all nodes in *G* are not matched **do**
    *M*\* = {}
    **for** *e* ∈ *E* sorted by best buddy weights **do**
      **if** *e* can be added to *M*\* **then**
        *M*\* = *M*\* ∪ *e*
      **end if**
    **end for**
    *M* = *M* ∪ *M*
    Remove nodes and edges which are part of *M*\* from G
**end while**
~~~

### 1.7 Iterative mis-join correction

Since the unitig layout is greedy, it can introduce errors by selecting a false Hi-C link which was not eliminated by our ratio scoring. These errors propagate downstream, causing large chimeric scaffolds and chromosomal fusions. We examine each join made within all the scaffolds in the last iteration for correctness. Any join with low spanning Hi-C support relative to the rest of the scaffold is broken and the links are blacklisted for further iterations.

We compute the physical coverage spanned by all read pairs aligned in a window of size *w* around each join. For each window, *w*, we create an auxiliary array, which stores 1 at position *i* if the physical coverage is greater than some cutoff *δ* and 1, otherwise. We then find the maximum sum subarray in this auxiliary array, since it captures the longest stretch of low physical coverage. If the position being tested for a mis-join lies within the region spanned by the maximal clique generated with the maximum sum subarray intervals for different cutoffs (Figure 2), the join is marked as incorrect. The physical coverage can be computed in *O*(*w* + *N*) time, where *N* is the number of read pairs aligned in window *w*. The maximum sum subarray computation takes *O*(*w*) time. If *K* is the number of cutoffs(*δ*) tested for the suspicious join finding, then the total runtime of mis-assembly detection becomes *O*(*K*(*N* + 2 * *w*)). The parameter *K* controls the specificity of the mis-assembly detection, thereby avoiding false positives. The algorithm for mis-join detection is sketched in Algorithm 2. When the majority of joins made in a particular iteration are flagged as incorrect by the algorithm, SASLA2 stops scaffolding and reports the scaffolds generated in the penultimate iteration as the final result.

#### Algorithm 2 Misjoin detection and correction algorithm

~~~
  *Cov* : Physical coverage array for a window size *w* around a scaffold join at position *p*
  on a scaffold
  *A* : Auxiliary array
  *I* : Maximum sum subarray intervals
  **for** δ ∈ {min_coverage, max_coverage} **do**
    **if** *Cov*[*i*] ≤ *δ* **then**
       *A*[*i*] = 1
    **else**
       *A*[*i*] = −1
    **end if**
    *s_δ_*, *e_δ_* = maximum_sum_subarray(*A*)
    *I* = *I* ∪ {*s_δ_*, *e_δ_*}
**end for**
*s*, *e* =maximal_clique_interval(*I*)
**if** *p* ∈ {*s*, *e*} **then**
    Break the scaffold at position *p*
**end if**
~~~

## 2 Results

### 2.1 Dataset description

We created artificial assemblies, each containing unitigs of same size, by splitting the GRCh38 [33] reference into fixed sized unitigs of 200 to 900 kbp. This gave us eight assemblies. The assembly graph for each input is built by adding edges for any adjacent unitigs in the genome.

For real data, we use the recently published NA12878 human dataset sequenced with Oxford Nanopore [34] and assembled with Canu [28]. We use a Hi-C library from Arima Genomics (Arima Genomics, San Diego, CA) sequenced to 40x Illumina coverage (SRX3651893). We compare results with the original SALSA(commit - 833fb11), SALSA2 with and without the assembly graph input(commit - 68a65b4), and 3D-DNA (commit - 3f18163). We did not compare our results with LACHESIS because it is no longer supported and is outperformed by 3D-DNA [20]. SALSA2 was run using default parameters, with the exception of graph incorporation, as listed. For 3D-DNA, alignments were generated using the Juicer alignment pipeline [35] with defaults (-m haploid -t 15000 -s 2), except for mis-assembly detection, as listed. A genome size of 3.2 Gbp was used for contiguity statistics for all assemblies.

For evaluation, we also used the GRCh38 reference to define a set of true and false links from the Hi-C graph. We mapped the assembly to the reference with MUMmer3.23 (nucmer -c 500 -l 20) [36] and generated a tiling using MUMmer’s show-tiling utility. For this “true link” dataset, any link joining unitigs in the same chromosome in the correct orientation was marked as true. This also gives the true unitig position, orientation, and chromosome assignment. We masked sequences in GRCh38 which matched known structural variants from a previous assembly of NA12878 [37] to avoid counting true variations as scaffolding errors.

### 2.2 Evaluation on simulated unitigs

#### 2.2.1 Assembly correction

We simulated assembly error by randomly joining 200 pairs of unitigs from each simulated assembly. All erroneous joins were made between unitigs that are more than 10 Mbp apart or were assigned to different chromosomes in the reference. The remaining unitigs were unaltered. We then aligned the Arima-HiC data and ran our assembly correction algorithm. When the algorithm marked a mis-join within 20 kbp of a true error we called it a true positive, otherwise we called it a false positive. Any unmarked error was called a false negative. The average sensitivity over all simulated assemblies was 77.62% and the specificity was 86.13%. The sensitivity was highest for larger unitigs (50% for 200 kbp versus more than 90% for untigs greater than 500 kbp, Supplementary Table S1) implying that our algorithm is able to accurately identify errors in large unitigs, which can have a negative impact on the final scaffolds if not corrected. Although we used a cutoff of 20 kbp to evaluate sensitivity and specificity, most of the predicted locations of misassembly were within 5 kbp from the true misassembly location (Supplementary Figure S2).

#### 2.2.2 Scaffold mis-join validation

As before, we simulated erroneous scaffolds by joining unitigs which were not within 10 Mbp in the reference or were assigned to different chromosomes. Rather than pairs of unitigs, each erroneous scaffold joined 10 unitigs and we generated 200 such erroneous scaffolds. The remaining unitigs were correctly scaffolded (ten unitigs per scaffold) based on their location in the reference. The average sensitivity was 67.66% and specificity was 100% (Supplementary Table S2)(no correct scaffolds were broken). Most of the un-flagged joins occurred near the ends of scaffolds and could be captured by decreasing the window size. Similar to assembly correction, we observed that sensitivity was highest with larger input unitigs. Most of the misjoins missed by the algorithm were near the ends of scaffolds. The issue in detecting mis-assemblies in these regions is the low Hi-C physical coverage. Also, the other missed joins were between the small contigs which are hard to capture with Hi-C data alone. This evaluation highlights the accuracy of the mis-join detection algorithm to avoid over-scaffolding and provide a suitable stopping condition.

#### 2.2.3 Scaffold accuracy

We evaluated scaffolds across three categories of error: orientation, order, and chimera. An orientation error occurs whenever the orientation of a unitig in a scaffold differs from that of the scaffold in the reference. An ordering error occurs when a set of three unitigs adjacent in a scaffold have non-monotonic coordinates in the reference. A chimera error occurs when any pair of unitigs adjacent in a scaffold align to different chromosomes in the reference. We broke the assembly at these errors and computed corrected scaffold lengths and NGA50 (analogous to the corrected NG50 metric defined by Salzberg et al. [38]). This statistic corrects for large but erroneous scaffolds which have an artificially high NG50. We did not include SALSA1 in the comparison because for small contig sizes (200 kbp to 500 kbp), none of the scaffolds contained more than 2 contigs. For larger sizes (600 kbp to 900 kbp), the contiguity widely varied depending upon the minimum confidence parameter for accepting links between contigs.

Hi-C scaffolding errors, particularly orientation errors, increased with decreasing assembly contiguity. We evaluated scaffolding methods across a variety of simulated unitig sizes. Figure 3 shows the comparison of these errors for 3D-DNA, SALSA2 without the assembly graph, and SALSA2 with the graph. SALSA2 produced fewer errors than 3D-DNA across all error types and input sizes. The number of correctly oriented unitigs increased significantly when assembly graph information was integrated with the scaffolding, particularly for lower input unitig sizes (Figure 3). For example, at 400 kbp, the orientation errors with the graph were comparable to the orientation errors of the graph-less approach at 900 kbp. The NGA50 for SALSA2 also increased when assembly graph information was included (Figure 4). This highlights the power of the assembly graph to improve scaffolding and correct errors, especially on lower contiguity assemblies. This also indicates that generating a conservative assembly, rather than maximizing contiguity, can be preferable for input to Hi-C scaffolding. All the assemblies described in this paper are available online and can be found at https://obj.umiacs.umd.edu/paper_assemblies/index.html.

**Fig 3.**
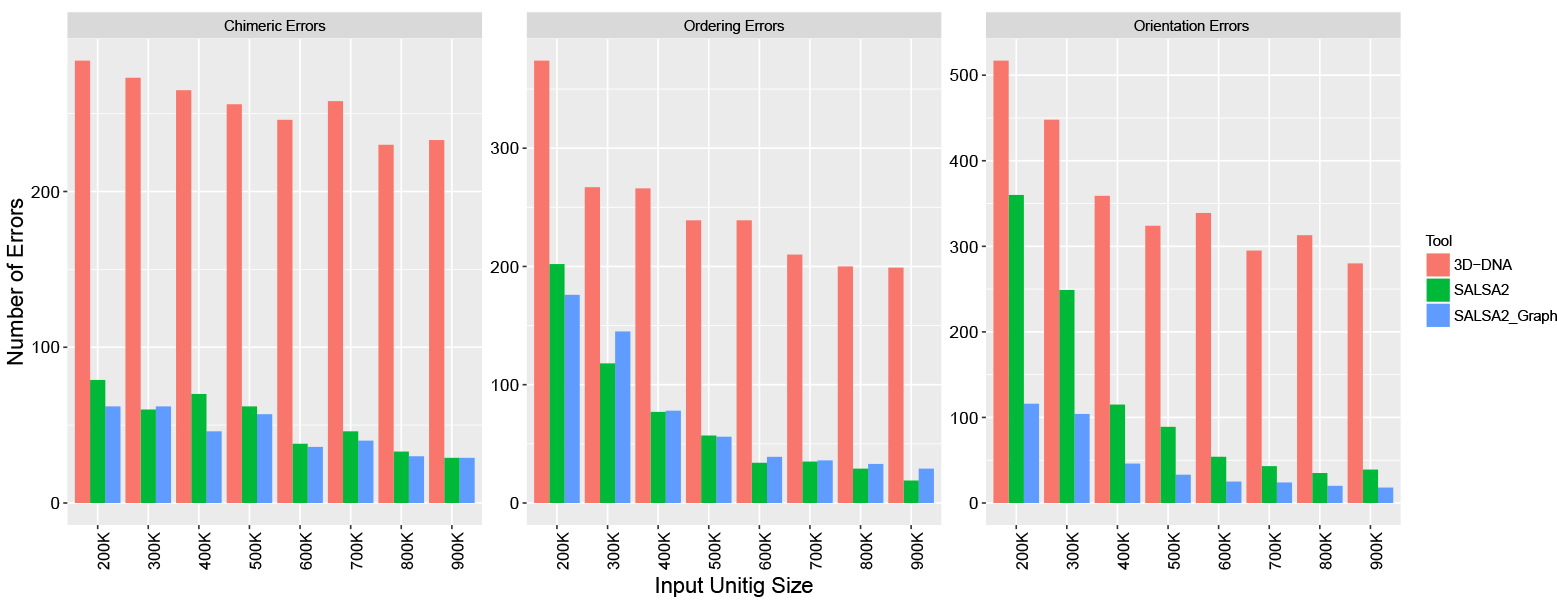
Comparison of orientation, ordering, and chimeric errors in the scaffolds produced by SALSA2 and 3D-DNA on the simulated data. As expected, the number of errors for all error types decrease with increasing input unitig size. Incorporating the assembly graph reduces error across all categories and most assembly sizes, with the largest decrease seen in orientation errors. SALSA2 utilizing the graph has 2-4 fold fewer errors than 3D-DNA.

**Fig 4.**
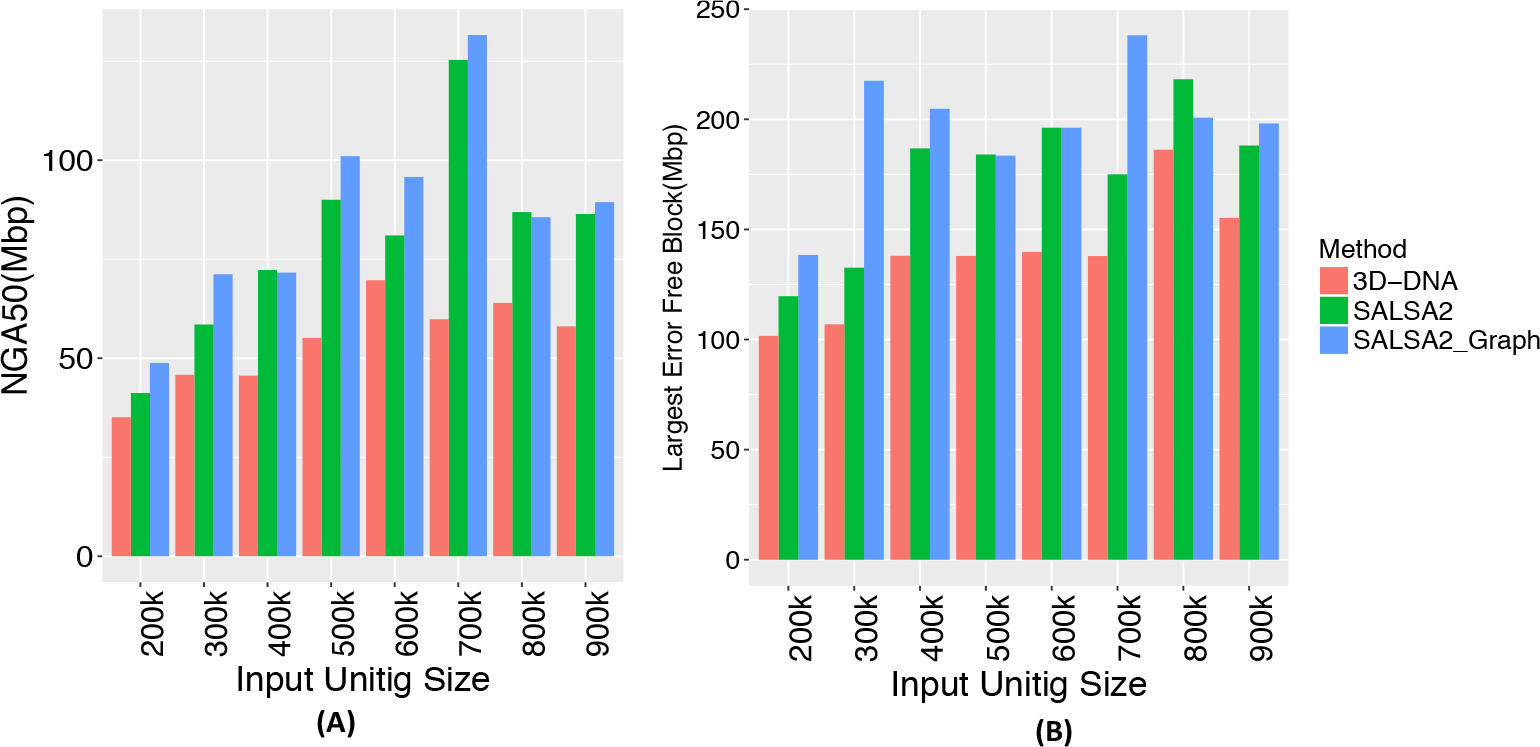
(A) NGA50 statistic for different input unitig sizes and (B) The length of longest error-free block for different input unitig sizes. Once again, the assembly graph typically increases both the NGA50 and the largest correct block.

### 2.3 Evaluation on NA12878

Table 1 lists the metrics for NA12878 scaffolds. We include an idealized scenario, using only reference-filtered Hi-C edges for comparison. As expected, the scaffolds generated using only true links had the highest NGA50 value and longest error-free scaffold block. SALSA2 scaffolds were generally more accurate and contiguous than the scaffolds generated by SALSA1 and 3D-DNA, even without use of the assembly graph. The addition of the graph further improved the NGA50 and longest error-free scaffold length.

**Table 1.** Scaffold and correctness statistics for NA12878 assemblies scaffolded with different Hi-C libraries. “True links” is an idealized case where the Hi-C links have been filtered in advance. The NG50 of human reference GRCh38 is 145 Mbp. The ratio between NG50 and NGA50 represents how many erroneous joins affect large scaffolds in the assembly. The bigger the difference between these values, the more aggressive the scaffolding was at the expense of accuracy. Longest chunk represents the longest error-free portion of the scaffolds. We observed that the 3D-DNA mis-assembly detection was overly aggressive in some cases, and so we ran some assemblies both with and without this feature. For the Illumina assembly as an input, 3D-DNA w correction did not finish within two weeks and is omitted. An evaluation of a previously published [20] 3D-DNA assembly from short-read contigs is included in Supplementary Table S3 but did not exceed SALSA2’s NGA50.

| Dataset | Method | NG50(Mbp) | NGA50(Mbp) | Longest Chunk (Mbp) | Orientation Errors | Ordering Errors | Chimeric Errors |
| --- | --- | --- | --- | --- | --- | --- | --- |
| Arima-HiC | SALSA2 true links | 83.31 | 79.48 | 172.19 | 78 | 101 | 0 |
|  | SALSA2 w graph | 112.08 | **71.54** | **164.46** | 102 | **106** | **90** |
|  | SALSA2 wo graph | **118.42** | 58.81 | 155.68 | 148 | 112 | 135 |
|  | 3D-DNA | 90.15 | 22.44 | 89.46 | 182 | 133 | 115 |
|  | SALSA1 | 19.09 | 14.81 | 73.14 | **99** | 176 | 96 |
| Mitotic Hi-C | SALSA2 w graph | 61.97 | **24.18** | **145.53** | **81** | **54** | **19** |
|  | SALSA1 | 27.88 | 15.62 | 85.71 | 142 | 78 | 120 |
|  | 3D-DNA w correction | 0.199 | 0.627 | 2.24 | 9775 | 10563 | 6159 |
|  | 3D-DNA wo correction | **85.56** | 17.18 | 70.18 | 250 | 215 | 164 |
| Chicago | SALSA2 w graph | 5.80 | 4.54 | 34.60 | **46** | 60 | **98** |
|  | SALSA1 | 5.21 | 3.94 | 34.60 | 83 | **21** | 187 |
|  | 3D-DNA w correction | 3.63 | 2.74 | 18.62 | 63 | 69 | 324 |
|  | 3D-DNA wo correction | **9.61** | **4.76** | **44.48** | 67 | 63 | 137 |
| Illumina Assembly | SALSA2 w graph | 96.78 | **7.99** | **43.56** | **1830** | **2299** | **635** |
|  | SALSA2 wo graph | 119.57 | 4.16 | 26.22 | 2225 | 2353 | 738 |
|  | 3D-DNA w correction | NA | NA | NA | NA | NA | NA |
|  | 3D-DNA wo correction | **176.09** | 1.00 | 13.12 | 5935 | 3433 | 2119 |

We also evaluated the assemblies using Feature Response Curves (FRC) based on scaffolding errors [40]. An assembly can have a high raw error count but still be of high quality if the errors are restricted to only short scaffolds. FRC captures this by showing how quickly error is accumulated, starting from the largest scaffolds. Figure 5(D) shows the FRC for different assemblies, where the X-axis denotes the cumulative % of assembly errors and the Y-axis denotes the cumulative assembly size. The assemblies with more area under the curve accumulate fewer errors in larger scaffolds and hence are more accurate. SALSA2 scaffolds with and without the graph have similar areas under the curve and closely match the curve of the assembly using only true links. The 3D-DNA scaffolds have the lowest area under the curve, implying that most errors in the assembly occur in the long scaffolds. This is confirmed by the lower NGA50 value for the 3D-DNA assembly (Table 1).

**Fig 5.**
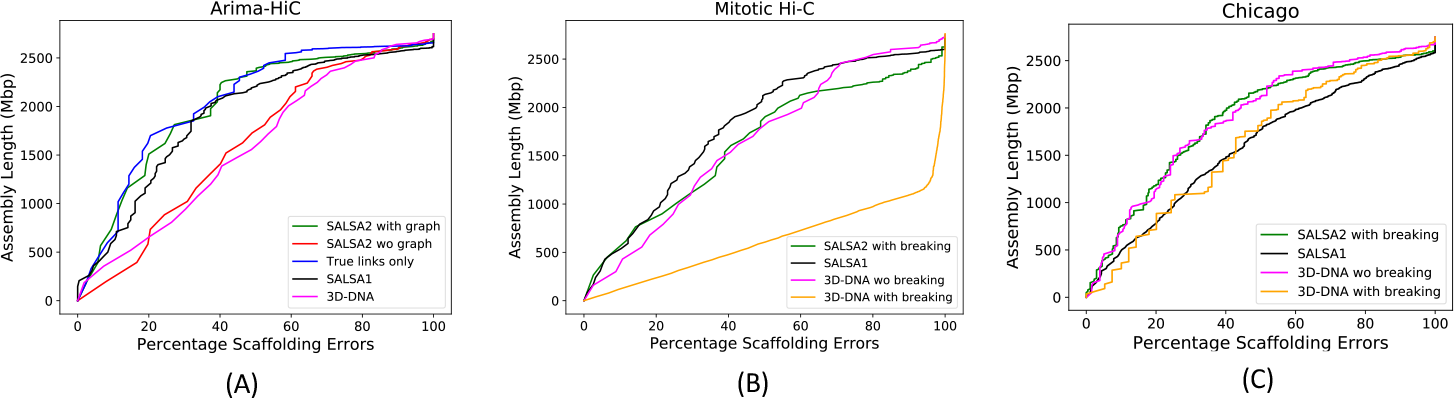
Feature Response Curve for (A) assemblies obtained from unitigs as input (B) assemblies obtained from mitotic Hi-C data and (C) assemblies obtained using Dovetail Chicago data. The best assemblies lie near the top left of the plot, with the largest area under the curve.

Apart from the correctness, SALSA2 scaffolds were highly contiguous and reached an NG50 of 112.8 Mbp (cf. GRCh38 NG50 of 145 Mbp). Figure 7 shows the alignment ideogram for the input unitigs as well as the SALSA2 assembly. Every color change indicates an alignment break, either due to error or due to the end of a sequence. The input unitigs are fragmented with multiple unitigs aligning to the same chromosome, while the SALSA2 scaffolds are highly contiguous and span entire chromosomes in many cases. Figure 6(A) shows the contiguity plot with corrected NG stats. As expected, the assembly generated with only true links has the highest values for all NGA stats. The curve for SALSA2 assemblies with and without the assembly graph closely matches this curve, implying that the scaffolds generated with SALSA2 are approaching the optimal assembly of this Arima-HiC data.

**Fig 6.**
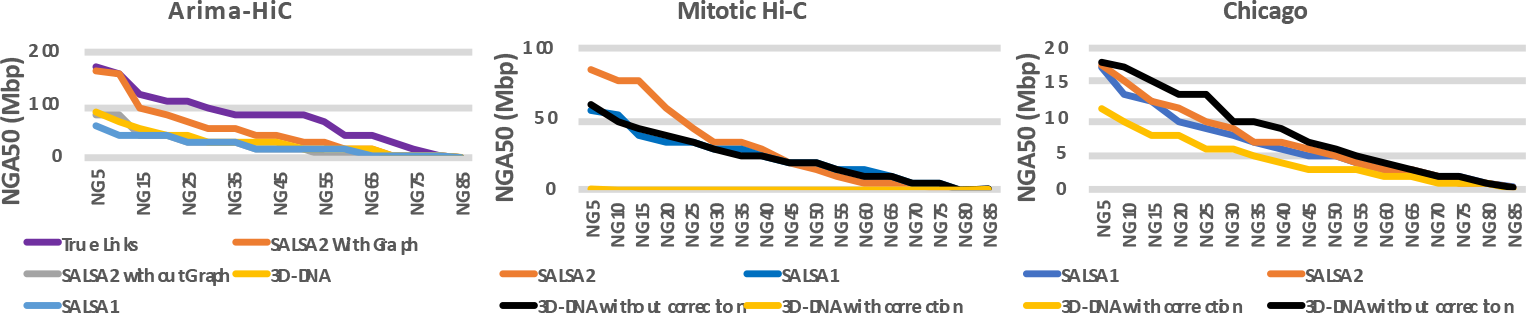
Contiguity plot for scaffolds generated with (A) standard Arima-HiC data (B) mitotic Hi-C data and (C) Chicago data. The X-axis denotes the NGAX statistic and the Y-axis denotes the corrected block length to reach the NGAX value. SALSA2 results were generated using the assembly graph, unless otherwise noted.

**Fig 7.**
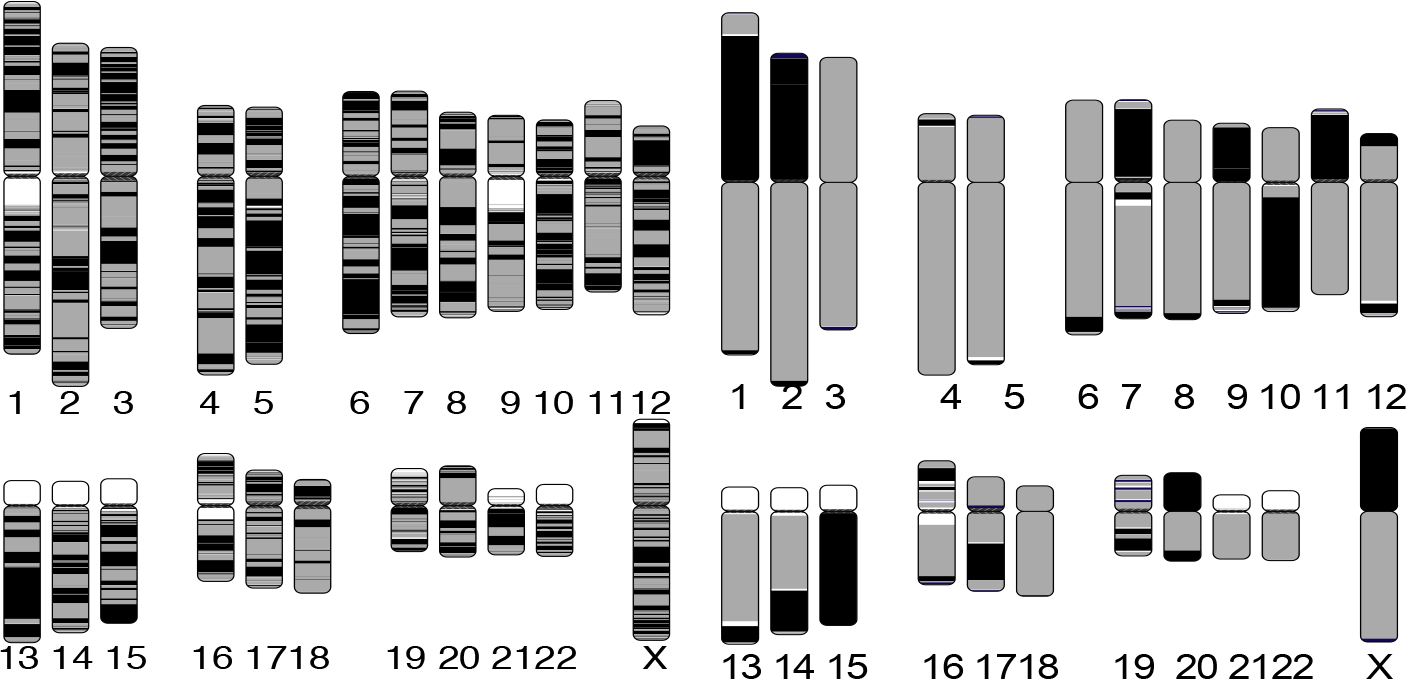
Chromosome ideogram generated using the coloredChromosomes [39] package. Each color switch denotes a change in the aligned sequence, either due to large structural error or the end of a unitig/scaffold. Left: input unitigs aligned to the GRCh38 reference genome. Right: SALSA2 scaffolds aligned to the GRCh38 reference genome. More than ten chromosomes are in a single scaffold. Chromosomes 16 and 19 are more fragmented due to scaffolding errors which break the alignment.

We also evaluated the ability of scaffolding short read assemblies for both 3D-DNA and SALSA2. We did not include SALSA1 in this comparison because it is not designed to scaffold short read assemblies. We observed that use of the assembly graph when scaffolding significantly reduced the number of orientation errors for SALSA2, increasing the scaffold NGA50 and largest chunk almost two-fold. When compared to 3D-DNA without input assembly correction, SALSA2 with the assembly graph generates scaffolds of much higher NGA50 (7.99 Mbp vs. 1.00 Mbp). The number of scaffolding errors across all the categories was much lower in SALSA2 compared to 3D-DNA.

We computed the CPU runtime and memory usage for both the methods while scaffolding long and short read assemblies with Arima-HiC data. We excluded the time required to map reads in both cases. While scaffolding the long-read assembly SALSA2 was 30-fold faster and required 3-fold less memory than 3D-DNA (11.44 CPU hours and 21.43 Gb peak memory versus 3D-DNA with assembly correction at 318 CPU hours and 64.66 Gb peak memory). For the short-read assembly, the difference in runtime was even more prounced. SALSA2 required 36.8 CPU hours and 61.8 Gb peak memory compared to 2980 CPU hours and 48.04 Gb peak memory needed by 3D-DNA without assembly correction. When run with assembly correction, 3D-DNA ran over 14 days on a 16-core machine without completing so we could not report any results.

### 2.4 Robustness to input library

We next tested scaffolding using two libraries with different Hi-C contact patterns. The first, from [42], is sequenced during mitosis. This removes the topological domains and generates fewer off-diagonal interactions. The other library was from [43], are *in vitro* chromatin sequencing library (Chicago) generated by Dovetail Genomics (L1). It also removes off-diagonal matches but has shorter-range interactions, limited by the size of the input molecules. As seen from the contact map in Figure 8, both the mitotic Hi-C and Chicago libraries follow different interaction distributions than the standard Hi-C (Arima-HiC in this case). We ran SALSA2 with defaults and 3D-DNA with both the assembly correction turned on and off.

**Fig 8.**
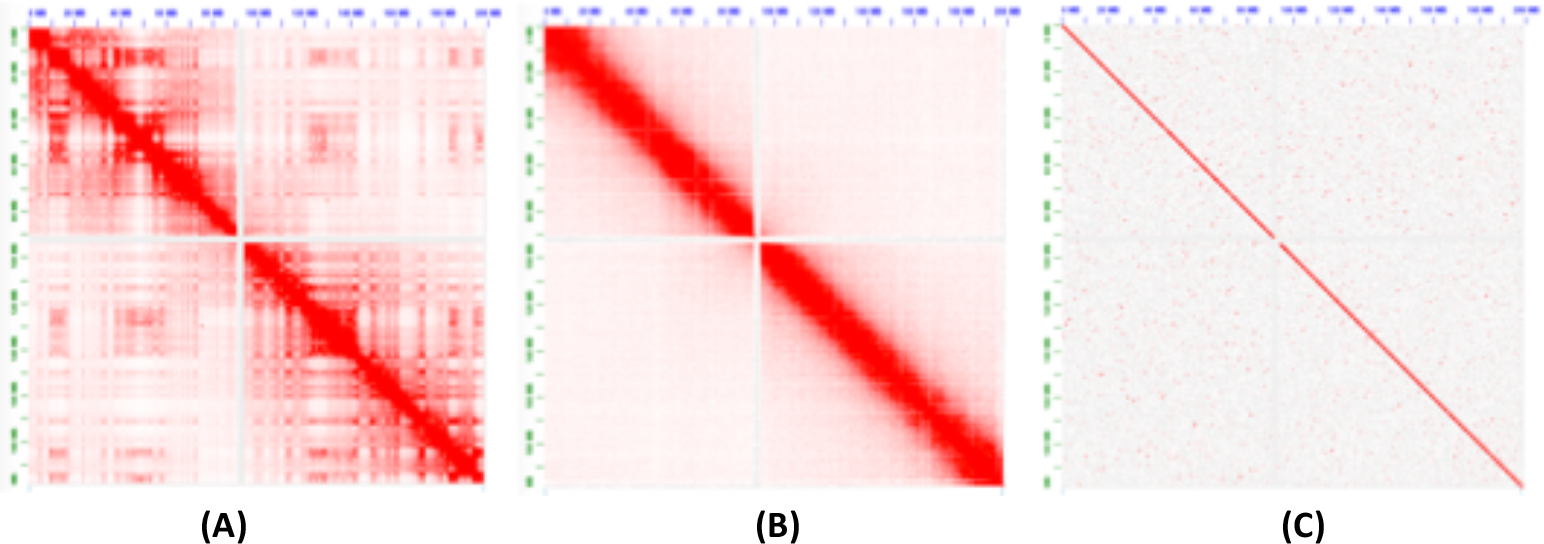
Contact map of Hi-C interactions on Chromosome 3 generated by the Juicebox software [41]. The cells sequenced in (A) normal conditions, (B) during mitosis, and (C) Dovetail Chicago

For mitotic Hi-C data, we observed that the 3D-DNA mis-assembly correction algorithm sheared the input assembly into small pieces, which resulted in more than 25,000 errors and more than half of the unitigs incorrectly oriented or ordered. Without mis-assembly correction, the 3D-DNA assembly has a higher number of orientation (250 vs. 81) and ordering (215 vs. 54) errors compared to SALSA2. The feature response curve for the 3D-DNA assembly with breaking is almost a diagonal (Figure 5(B)) because the sheared unitigs appeared to be randomly joined. SALSA2 scaffolds contain longer stretches of correct scaffolds compared to 3D-DNA with and without mis-assembly correction (Figure 6(B)). SALSA1 scaffolds had a similar error count to SALSA2 but were less contiguous.

For the Chicago libraries, 3D-DNA without correction had the best NGA50 and largest correct chunk. However, the scaffolds had more chimeric join errors than SALSA2. SALSA2 outperformed 3D-DNA in terms of NG50, NGA50, and longest chunk when 3D-DNA was run with assembly correction. 3D-DNA uses signatures of chromosome ends [20] to identify break positions which are not prominently present in Chicago data. As a result, it generated more chimeric joins compared to SALSA2.

However, the number of order and orientation errors was similar across the methods. Since Chicago libraries do not provide chromosome-spanning contact information for scaffolding, the NG50 value for SALSA2 is 5.8 Mbp, comparable to the equivalent coverage assembly (50% L1+L2) in [43] but much smaller than Hi-C libraries.

Interestingly, SALSA1 was able to generate scaffolds of similar contiguity to SALSA2, which can be attributed to the lack of long range contact information in the library. SALSA2 is robust to changing contact distributions. In the case of Chicago data it produced a less contiguous assembly due to the shorter interaction distance. However, it avoids introducing false chromosome joins, unlike 3D-DNA, which appears tuned for a specific contact model.

## 3 Conclusion

In this work, we present the first Hi-C scaffolding method that integrates an assembly graph to produce high-accuracy, chromosome-scale assemblies. Our experiments on both simulated and real sequencing data for the human genome demonstrate the benefits of using an assembly graph to guide scaffolding. We also show that SALSA2 outperforms alternative Hi-C scaffolding tools on assemblies of varied contiguity, using multiple Hi-C library preparations.

SALSA2’s misassembly correction and scaffold misjoin validation can be improved in several ways. The current implementation doesn’t detect a misjoin between two small contigs with high accuracy, mainly because Hi-C data doesn’t have enough resolution to correct such errors. Also, we don’t account for any GC bias correction when using the Hi-C coverage for detecting misjoins. Addressing these challenges in misjoin detection and misassembly correction is the immediate next step to improve the SALSA2 software.

Hi-C scaffolding has been historically prone to inversion errors when the input assembly is highly fragmented. The integration of the assembly graph with the scaffolding process can overcome this limitation. Orientation errors introduced in the assembly and scaffolding process can lead to incorrect identification of structural variations. On simulated data, more than 50% of errors were due to inversions, and integrating the assembly graph reduced these by as much as 3 to 4 fold. We did not observe as much improvement with the NA12878 test dataset because the contig NG50 was much higher than in the simulation. However, it is not always possible to assemble multi-megabase contigs. In such cases, the assembly graph is useful for limiting Hi-C errors.

Most existing Hi-C scaffolding methods also require an estimate for the number of chromosomes for a genome. This is implicitly taken to be the desired number of scaffolds to output. As demonstrated by the Chicago, mitotic, and replicate [44] Hi-C libraries, the library as well as the genome influences the maximum correct scaffold size. It can be impractical to sweep over hundreds of chromosome values to select a “best” assembly. Since SALSA2’s mis-join correction algorithm stops scaffolding after the useful linking information in a dataset is exhausted, no chromosome count is needed as input.

Obtaining the chromosome-scale picture of the genome is important and there is a trade-off between accuracy and continuity of the assembly. However, we believe that manual curation to remove assembly errors is an expensive and involved process which can often outpace the cost of the rest of the project. Most of the assembly projects using Hi-C data have had a significant curation effort to date [19, 45]. Thus, we believe that not introducing errors in the first place is a better strategy to avoid the later burden of manual curation of small errors in chromosomes. The Hi-C data can be used with other independent technologies, such as optical mapping or linked-reads to reach accurate chromosome-scale scaffolds. 3D-DNA was recently updated to not require the chromosome count as input but the algorithm used has not been described.

Interestingly, it no longer generates single chromosome scaffolds in our experiments, a major claim in [20], supporting a conservative scaffolding approach. Even while scaffolding short-read assemblies, we observed that SALSA2 generated more accurate scaffolds than 3D-DNA, indicating the utility of SALSA2 in scaffolding existing short-read assemblies of different genomes with the newly generated Hi-C data.

As the Genome10K consortium [46] and independent scientists begin to sequence novel lineages in the tree of life, it may be impractical to generate physical or genetics maps for every organism. Thus, Hi-C sequencing combined with SALSA2 presents an economical alternative for the reconstruction of chromosome-scale assemblies

## Acknowledgments

AS and SS were funded by generous support from NHGRI (grant# 1R44HG009584). JG and MP were supported by NIH grant R01-AI-100947 to MP. SK, AR, BPW, and AMP were supported by the Intramural Research Program of the National Human Genome Research Institute, National Institutes of Health. AR was also supported by a grant from the Korean Visiting Scientist Training Award (KVSTA) through the Korea Health Industry Development Institute (KHIDI), funded by the Ministry of Health & Welfare, Republic of Korea (grant number: HI17C2098). This work utilized the computational resources of the NIH HPC Biowulf cluster (http://hpc.nih.gov).

## References

1. Nagarajan N, Pop M. Sequence assembly demystified. Nature Reviews Genetics. 2013;14(3):157–167.

2. Miller JR, Koren S, Sutton G. Assembly algorithms for next-generation sequencing data. Genomics. 2010;95(6):315–327.

3. Pevzner PA, Tang H, Waterman MS. An Eulerian path approach to DNA fragment assembly. Proceedings of the National Academy of Sciences. 2001;98(17):9748–9753.

4. Myers EW. The fragment assembly string graph. Bioinformatics. 2005;21(suppl 2):ii79–ii85.

5. Nagarajan N, Pop M. Parametric complexity of sequence assembly: theory and applications to next generation sequencing. Journal of computational biology. 2009;16(7):897–908.

6. Venter JC, Smith HO, Hood L. A new strategy for genome sequencing. Nature. 1996;381(6581):364.

7. Gnerre S, MacCallum I, Przybylski D, Ribeiro FJ, Burton JN, Walker BJ, et al. High-quality draft assemblies of mammalian genomes from massively parallel sequence data. Proceedings of the National Academy of Sciences. 2011;108(4):1513–1518.

8. Schwartz DC, Li X, Hernandez LI, Ramnarain SP, Huff EJ, Wang YK. Ordered restriction maps of Saccharomyces cerevisiae chromosomes constructed by optical mapping. Science. 1993;262(5130):110–114.

9. Dong Y, Xie M, Jiang Y, Xiao N, Du X, Zhang W, et al. Sequencing and automated whole-genome optical mapping of the genome of a domestic goat (Capra hircus). Nature biotechnology. 2013;31(2):135–141.

10. Shelton JM, Coleman MC, Herndon N, Lu N, Lam ET, Anantharaman T, et al. Tools and pipelines for BioNano data: molecule assembly pipeline and FASTA super scaffolding tool. BMC genomics. 2015;16(1):734.

11. Zheng GX, Lau BT, Schnall-Levin M, Jarosz M, Bell JM, Hindson CM, et al. Haplotyping germline and cancer genomes with high-throughput linked-read sequencing. Nature biotechnology. 2016;.

12. Weisenfeld NI, Kumar V, Shah P, Church DM, Jaffe DB. Direct determination of diploid genome sequences. Genome research. 2017;27(5):757–767.

13. Yeo S, Coombe L, Warren RL, Chu J, Birol I. Arcs: Scaffolding genome drafts with linked reads. Bioinformatics. 2017;.

14. Simonis M, Klous P, Splinter E, Moshkin Y, Willemsen R, De Wit E, et al. Nuclear organization of active and inactive chromatin domains uncovered by chromosome conformation capture–on-chip (4C). Nature genetics. 2006;38(11):1348–1354.

15. Lieberman-Aiden E, Van Berkum NL, Williams L, Imakaev M, Ragoczy T, Telling A, et al. Comprehensive mapping of long-range interactions reveals folding principles of the human genome. science. 2009;326(5950):289–293.

16. Burton JN, Adey A, et al. Chromosome-scale scaffolding of de novo genome assemblies based on chromatin interactions. Nature biotechnology. 2013;31(12):1119–1125.

17. Kaplan N, Dekker J. High-throughput genome scaffolding from in vivo DNA interaction frequency. Nature biotechnology. 2013;31(12):1143–1147.

18. Marie-Nelly H, Marbouty M, Cournac A, Flot JF, Liti G, Parodi DP, et al. High-quality genome (re) assembly using chromosomal contact data. Nature communications. 2014;5.

19. Bickhart DM, Rosen BD, Koren S, et al. Single-molecule sequencing and chromatin conformation capture enable de novo reference assembly of the domestic goat genome. Nature Genetics. 2017;49(4):643–650.

20. Dudchenko O, Batra SS, Omer AD, et al. De novo assembly of the Aedes aegypti genome using Hi-C yields chromosome-length scaffolds. Science. 2017;356(6333):92–95.

21. Ghurye J, Pop M, Koren S, Bickhart D, Chin CS. Scaffolding of long read assemblies using long range contact information. BMC genomics. 2017;18(1):527.

22. Dixon JR, Selvaraj S, Yue F, Kim A, Li Y, Shen Y, et al. Topological domains in mammalian genomes identified by analysis of chromatin interactions. Nature. 2012;485(7398):376–380.

23. Eid J, Fehr A, Gray J, Luong K, Lyle J, Otto G, et al. Real-time DNA sequencing from single polymerase molecules. Science. 2009;323(5910):133–138.

24. Jain M, Olsen HE, Paten B, Akeson M. The Oxford Nanopore MinION: delivery of nanopore sequencing to the genomics community. Genome biology. 2016;17(1):239.

25. Li H. Minimap and miniasm: fast mapping and de novo assembly for noisy long sequences. Bioinformatics. 2016;32(14):2103–2110.

26. Li H, Durbin R. Fast and accurate short read alignment with Burrows–Wheeler transform. Bioinformatics. 2009;25(14):1754–1760.

27. Wysoker A, Tibbetts K, Fennell T. Picard tools version 1.90; 2013.

28. Koren S, Walenz BP, Berlin K, et al. Canu: scalable and accurate long-read assembly via adaptive k-mer weighting and repeat separation. Genome research. 2017;27(5):722–736.

29. Chin CS, Peluso P, Sedlazeck FJ, Nattestad M, Concepcion GT, Clum A, et al. Phased Diploid Genome Assembly with Single Molecule Real-Time Sequencing. bioRxiv. 2016;doi:10.1101/056887.

30. Kolmogorov M, Yuan J, Lin Y, Pevzner P. Assembly of Long Error-Prone Reads Using Repeat Graphs. bioRxiv. 2018;doi:10.1101/247148.

31. Edmonds J. Paths, trees, and flowers. Canadian Journal of mathematics. 1965;17(3):449–467.

32. Poloczek M, Szegedy M. Randomized greedy algorithms for the maximum matching problem with new analysis. In: Foundations of Computer Science (FOCS), 2012 IEEE 53rd Annual Symposium on. IEEE; 2012. p. 708–717.

33. Schneider VA, Graves-Lindsay T, Howe K, Bouk N, Chen HC, Kitts PA, et al. Evaluation of GRCh38 and de novo haploid genome assemblies demonstrates the enduring quality of the reference assembly. Genome research. 2017;27(5):849–864.

34. Jain M, Koren S, Quick J, Rand AC, et al. Nanopore sequencing and assembly of a human genome with ultra-long reads. bioRxiv. 2017; p. 128835.

35. Durand NC, Shamim MS, et al. Juicer provides a one-click system for analyzing loop-resolution Hi-C experiments. Cell systems. 2016;3(1):95–98.

36. Kurtz S, Phillippy A, Delcher AL, Smoot M, Shumway M, Antonescu C, et al. Versatile and open software for comparing large genomes. Genome biology. 2004;5(2):R12.

37. Pendleton M, Sebra R, Pang AWC, et al. Assembly and diploid architecture of an individual human genome via single-molecule technologies. Nature methods. 2015;.

38. Salzberg SL, Phillippy AM, Zimin A, Puiu D, Magoc T, Koren S, et al. GAGE: A critical evaluation of genome assemblies and assembly algorithms. Genome research. 2012;22(3):557–567.

39. Böhringer S, Gödde R, Böhringer D, Schulte T, Epplen JT. A software package for drawing ideograms automatically. Online J Bioinformatics. 2002;1:51–61.

40. Vezzi F, Narzisi G, Mishra B. Feature-by-feature–evaluating de novo sequence assembly. PloS one. 2012;7(2):e31002.

41. Durand NC, Robinson JT, Shamim MS, et al. Juicebox provides a visualization system for Hi-C contact maps with unlimited zoom. Cell systems. 2016;3(1):99–101.

42. Naumova N, Imakaev M, Fudenberg G, et al. Organization of the mitotic chromosome. Science. 2013;342(6161):948–953.

43. Putnam NH, O’Connell BL, Stites JC, Rice BJ, et al. Chromosome-scale shotgun assembly using an in vitro method for long-range linkage. Genome research. 2016;26(3):342–350.

44. Fletez-Brant K, Qiu Y, Gorkin DU, Hu M, Hansen KD. Removing unwanted variation between samples in Hi-C experiments. bioRxiv. 2017;doi:10.1101/214361.

45. Matthews BJ, Dudchenko O, Kingan S, Koren S, Antoshechkin I, Crawford JE, et al. Improved Aedes aegypti mosquito reference genome assembly enables biological discovery and vector control. bioRxiv. 2017; p. 240747.

46. Koepfli KP, Paten B, of Scientists GKC, O?Brien SJ. The Genome 10K Project: a way forward. Annu Rev Anim Biosci. 2015;3(1):57–111.

